# Double-strand breaks in facultative heterochromatin require specific movements and chromatin changes for efficient repair

**DOI:** 10.1101/2023.12.01.569547

**Authors:** Marieke R. Wensveen, Aditya A. Dixit, Serafin U. Colmenares, Aniek Janssen

## Abstract

DNA double-strand breaks (DSBs) must be properly repaired within diverse chromatin domains to maintain genome stability. Whereas euchromatin has an open structure and is associated with active transcription, facultative heterochromatin is essential to silence developmental genes and forms compact nuclear condensates, called polycomb bodies. Whether the specific chromatin properties of facultative heterochromatin require distinct DSB repair mechanisms remains unknown. Here, we integrate single DSB systems in euchromatin and facultative heterochromatin in *Drosophila melanogaster* and find that facultative heterochromatic DSBs rapidly move outside polycomb bodies. These DSB movements coincide with a break-proximal reduction in the canonical heterochromatin mark histone H3 Lysine 27 trimethylation (H3K27me3). We demonstrate that DSB movement and loss of H3K27me3 at heterochromatic DSBs both depend on the histone demethylase dUtx. Moreover, loss of dUtx specifically disrupts completion of homologous recombination at heterochromatic DSBs. We conclude that DSBs in facultative heterochromatin require dUtx-mediated loss of H3K27me3 to promote DSB movement and repair.

## INTRODUCTION

Eukaryotic cells are continuously exposed to factors that break or chemically alter DNA. One particularly dangerous type of DNA damage is a double-strand break (DSB), in which both strands of the DNA helix are severed. Improper repair of DSBs can directly lead to insertions, deletions and chromosomal rearrangements associated with disease development including cancer^1^. To overcome these detrimental outcomes, cells have evolved mechanisms to repair DSBs of which the two main pathways are Non-Homologous End Joining (NHEJ) and Homologous Recombination (HR). During NHEJ, the severed DNA ends undergo limited end processing and are directly ligated, which can result in small insertions or deletions (indels) at the break site and is therefore considered error-prone^2^. HR, on the other hand, is usually more precise since it relies on a homologous template to repair the DSB. During HR, the broken DNA ends undergo 5’ to 3’ end resection, generating 3’ single-stranded DNA (ssDNA) overhangs. This overhang invades a homologous sequence on the sister chromatid or the homologous chromosome, which serves as a repair template. The choice of DSB repair pathway depends on multiple aspects, including cell cycle phase, the sequence context of the surrounding DNA, as well as the pre-existing chromatin state^2–4^.

The eukaryotic nucleus consists of a variety of chromatin domains each characterized by specific molecular and biophysical properties. Whereas euchromatin has an open chromatin structure with actively transcribed genes, heterochromatin is more condensed and transcriptionally inactive. One type of heterochromatin is facultative heterochromatin, which is essential to silence specific developmental genes. Facultative heterochromatin can cover large genomic distances (e.g. developmental genes such as Hox genes)^5^, or regulatory regions (e.g. promoters)^6^. This type of heterochromatin is enriched for Histone H3 Lysine 27 trimethylation (H3K27me3) and polycomb group (PcG) proteins, and accumulates in nuclear foci, called polycomb bodies^7–9^. Polycomb bodies cluster PcG-bound transcriptionally repressed genomic regions to maintain correct silencing of developmental genes^9–13^. Although the DSB response in open, euchromatic regions has been extensively studied, the DSB repair response in facultative heterochromatin remains largely unknown.

In the past decade, it has become clear that the pre-existing chromatin state can directly influence the DSB repair response. For example, DSBs in actively transcribed regions are prone to clustering and repair by HR^14,15^. Moreover, DSBs in centromeres^16^, nucleoli^17–19^ and constitutive heterochromatin domains^16,20– 22^ have been found to move outside the respective domains to facilitate repair. Previous evidence suggests that DSB movements can also occur within the inactive X chromosome^23^, which is a specific type of facultative heterochromatin enriched for both H3K27me3 and H3K9me3^24^. Irradiation of female human fibroblasts resulted in the specific exclusion of DSB repair proteins outside the inactive X chromosome, suggestive of DSB movement^23^. Moreover, decompaction of the inactive X upon laser irradiation has also been observed^25^. Nevertheless, live imaging of individual DSBs to precisely monitor their dynamics within facultative heterochromatin has never been performed. More importantly, the response of polycomb bodies to DSBs in a physiological, *in vivo* setting, and whether this chromatin environment facilitates particular movements of DSBs, remains unknown.

Various histone modifications have been identified to play a role in DSB repair in euchromatin^3,26^. Silencing histone modifications, including H3K27me3^27,28^ and H3K9me2/3^29,30^ have been described to be deposited at DSBs in euchromatin, resulting in local, transient heterochromatinization and transcriptional silencing. To restart transcription after DSB repair in euchromatin, active removal of H3K27me3 by the mammalian histone demethylase UTX (Ubiquitously transcribed Tetratricopeptide repeat on X chromosome) has been suggested to occur specifically in cancer cells, not healthy fibroblasts^31^. In contrast to the accumulation of silencing marks at euchromatic DSBs, we previously identified a loss of H3K9me2/3 at DSBs within *Drosophila* constitutive heterochromatin^32,33^. These findings suggest that eu- and heterochromatin regions require differential changes in silencing histone modifications to repair their DSBs. Whether specific H3K27-modifying activities are needed to repair DSBs in H3K27me3-enriched facultative heterochromatin domains remains untested.

Here, we study the dynamic DSB response in facultative heterochromatin *in vivo* by integrating inducible single DSB systems^22,34^ in euchromatin and facultative heterochromatin regions in the fruit fly *Drosophila melanogaster*. Using high-resolution live imaging, we find that the majority of DSBs in polycomb bodies rapidly move outside these domains. Moreover, we find that facultative heterochromatic DSBs specifically undergo a local decrease in the canonical heterochromatin histone mark H3K27me3, which is mediated by the histone demethylase dUtx. Early steps of HR can occur efficiently within polycomb bodies and are independent of dUtx, while dUtx is required for subsequent DSB movement and completion of HR. Together, our results reveal that DSBs in facultative heterochromatin move outside the compacted polycomb bodies to promote timely repair by HR.

## RESULTS

### Development of a single double-strand break system in facultative heterochromatin

To study DSB repair in facultative heterochromatin in detail in animal tissue, and directly compare responses to euchromatin, we generated a set of inducible single DSB systems in *Drosophila*. We integrated our previously established *in vivo* DR-*white* reporters^22,34^ into three facultative heterochromatin regions and two euchromatin regions (**Fig.1A, B**). Upon expression of I-SceI, DSBs are induced in the upstream *white* gene. DSB repair pathway usage can subsequently be determined by sequencing the resulting repair products; HR with the downstream i*white* sequence will generate an intact upstream *white* gene, while NHEJ will generate small indels at the cut site (**Fig.1A**). We performed ChIP-qPCR (Chromatin Immuno-Precipitation followed by quantitative PCR (qPCR)) for the canonical facultative heterochromatin histone modification H3K27me3 and confirmed enrichment of H3K27me3 at the three DR-*white* integrations in heterochromatin when compared to the two euchromatic insertions (**Fig.1C**). Moreover, internal controls revealed strong specificity of the antibody used for H3K27me3 ChIP analysis (**Fig.S1A**).

**Figure 1.**
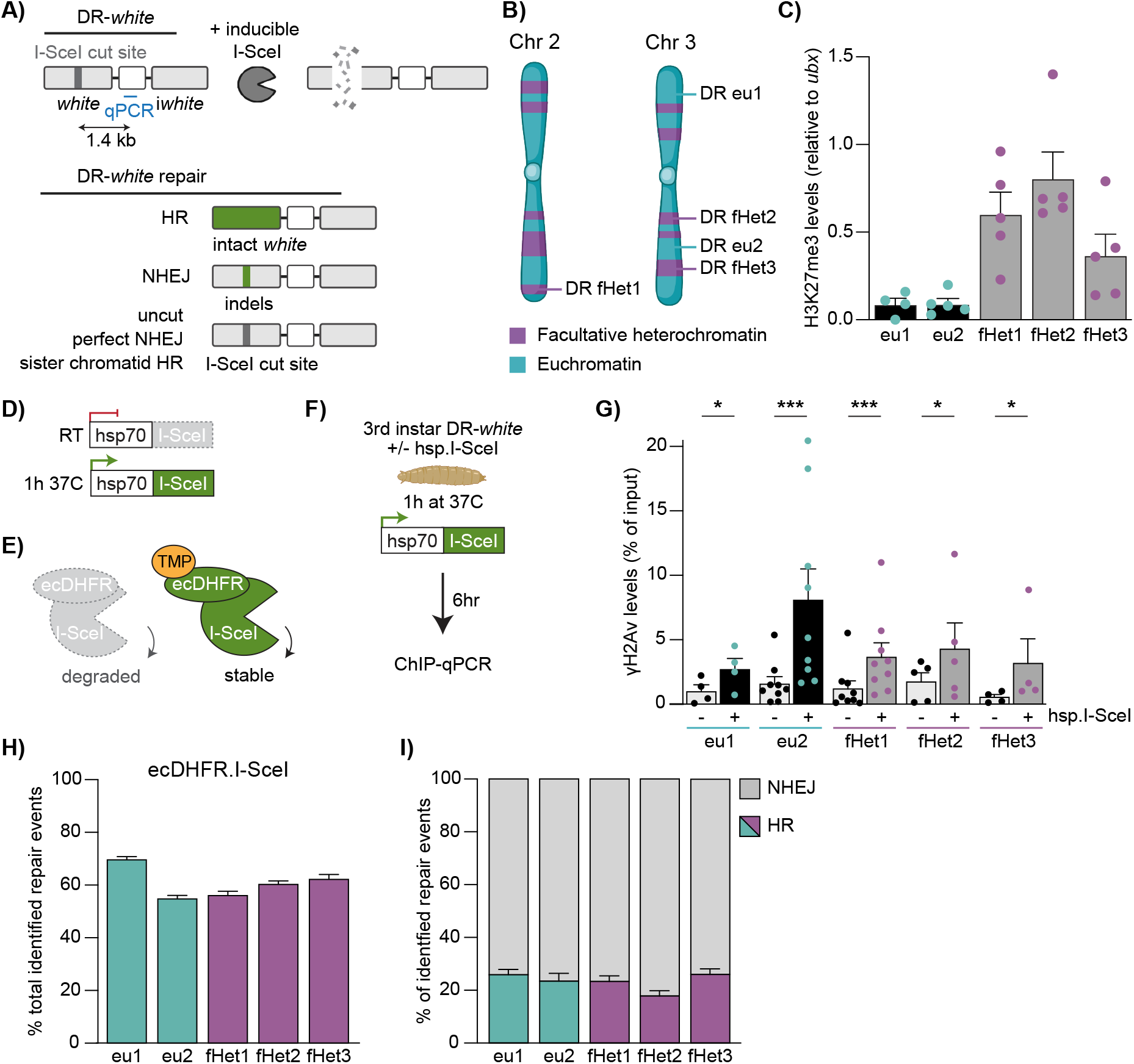
DR-*white* system to induce single DSBs in euchromatin and facultative heterochromatin. **A**) Schematic of the DR-*white* system. Inducible expression of I-SceI creates a single DSB in the upstream *white* gene. HR with the downstream (5’ and 3’) truncated i*white* sequence results in loss of the I-SceI cut site (-23bp), and thereby an intact upstream *white* gene. NHEJ can result in indels at the DSB site. Perfect NHEJ or HR using the sister chromatid can lead to recreation of the I-SceI cut site. **B**) Schematic of DR-*white* integration sites in the fly genome (2x in euchromatin [DR *eu1*, DR *eu2*] and 3x in facultative heterochromatin [DR *fHet1*, DR *fHet2*, DR *fHet3*]). **C**) ChIP-qPCR analysis for H3K27me3 at the DR-*white* locus using qPCR primers that bind 1.4kb downstream of the I-SceI cut site (indicated in Fig 1A). H3K27me3 levels were normalized using *ubx* gene qPCR primers as a positive internal control (*ubx* has consistently high H3K27me3 levels). Averages are shown for ≥4 independent experiments +SEM. **D**) Schematic of the hsp.I-SceI construct. The hsp70 promoter upstream of I-SceI can be activated by shiiing larvae for one hour to 37°C. RT = room temperature. **E**) Schematic of the ecDHFR-I-SceI system. Proteasomal degradation of ecDHFR-I-SceI can be blocked by adding the stabilizing compound trimethoprim (TMP). **F**) Experimental set up for ChIP-qPCR experiments upon DSB induction using hsp.I-SceI. 3^rd^ instar DR-*white* larvae with and without hsp.I-SceI (control) were heat-shocked for one hour at 37°C to activate the *hsp70* promoter. Six hours later chromatin was extracted from 40 larvae per condition and subjected to ChIP-qPCR using the qPCR primers indicated in Fig 1A. **G**) ChIP-qPCR analysis for γH2Av in the absence (-) and presence (+) of hsp.I-SceI (as in Fig 1F). **H, I**) DR-*white*/ecDHFR-I-SceI larvae were fed trimethoprim (TMP) for 3-4 days to induce I-SceI. Repair products at the upstream *white* gene were PCR amplified, Sanger sequenced and analyzed using the TIDE algorithm^36^. Graphs show quantification of total identified repair events in all PCR products (**H**) and the percentage of repair products with either HR (color) or NHEJ (grey) signatures (**I**). Bars indicate averages +SEM of ≥9 independent experiments (larvae) per condition. (*) p-value≤0.05, (***) p-value≤0.001, ratio paired t-test (**G**).

To allow timed DSB induction, we combined our DR-*white* systems with either a heat-shock inducible I-SceI transgene (*hsp70*.I-SceI, **Fig.1D**) or an ecDHFR-I-SceI transgene, which depends on the ligand trimethoprim to stabilize the ecDHFR-I-SceI protein (**Fig.1E**)^22^. We first tested the efficiency and inducibility of our DR-*white* systems by performing ChIP-qPCR for phosphorylated H2Av on Serine 137 (γH2Av, γH2AX in mammals), one of the earliest chromatin markers of DSB induction^35^. Heat-shock inducible expression of I-SceI resulted in a local increase in γH2Av levels within six hours at both euchromatic and heterochromatic DR-*white* loci (**Fig.1F, G**). Moreover, we find the appearance of single γH2Av foci in nuclei of imaginal discs six hours after heat-shock inducible I-SceI expression in larvae containing either a eu-or heterochromatic DR-*white* insertion (16-18% of cells contained a single γH2Av focus compared to 3-5% in control cells) (**Fig.S1B**). These results suggest efficient DSB induction in both eu- and heterochromatic loci.

To directly determine which DSB repair pathways play a role in facultative heterochromatin, we performed Sanger sequencing followed by TIDE analysis^36^ on the DR-*white* reporters upon I-SceI expression. Feeding trimethoprim throughout development (3-4 days) to DR-*white* larvae expressing ecDHFR-I-SceI (**Fig.1E**) results in the appearance of repair products in both eu- and heterochromatin (**Fig.1H**). Repair rates vary slightly between integration sites (55-70%), possibly reflecting differential cufng efficiency or repair timing at the different sites. However, no apparent differences in the number of identified repair products were found between heterochromatin and euchromatin, indicating that both domains undergo efficient DSB induction and repair. More importantly, repair pathway analyses revealed that I-SceI-induced DSBs in both eu- and heterochromatin regions employ HR (17-26%) and NHEJ (74-83%) to a similar extent (**Fig.1I**). In line with this, expression of hsp.I-SceI also resulted in the appearance of DSB repair products (7-19%) and similar HR and NHEJ repair percentages in eu- and heterochromatin DR-*white* integrations (**Fig.S1C, D**). Our heat-shock inducible system resulted in an overall lower number of repair products when compared to ecDHFR-I-SceI, likely reflecting the shorter duration of I-SceI expression (24 hours in hsp.I-SceI compared to 3-4 days in ecDHFR-I-SceI system). To confirm the role of HR repair at facultative heterochromatic DSBs, we depleted the end resection HR protein CtIP using RNAi (**Fig.S1E**) and analyzed DR-*white* repair products following hsp.I-SceI induction (**Fig.S1F, G**). As expected, loss of CtIP resulted in a decrease in HR repair products (3-11% compared to 23-33% in control) in both eu- and heterochromatin, indicating that the identified HR repair products indeed reflect end resection dependent HR repair.

Together, these data reveal that our systems efficiently induce DSBs with little variation between euchromatin and facultative heterochromatin regions in repair products, suggesting that both chromatin regions undergo similar DSB repair efficiency and pathway choice. Our inducible single DSB system *in vivo* therefore allows us to perform detailed analyses of DSB repair in facultative heterochromatin, and directly compare it to the DSB response in euchromatin.

### DSBs rapidly move outside polycomb bodies

Specific DSB spatiotemporal dynamics are associated with a variety of chromatin domains, such as centromeres^16^, nucleoli^17–19^ and constitutive heterochromatin^16,20,22^. These dynamics include the movement of DSBs to the periphery of the respective domain^19,21^. Facultative heterochromatin forms distinct domains in the fly and mammalian nucleus, termed polycomb bodies^7–9^. We wished to determine whether the distinct molecular- and biophysical-properties of polycomb bodies^37–40^ could impact DSB dynamics and promote movements similar to those previously identified in other nuclear domains. To this end, we employed our DR-*white* systems to perform *in vivo* live imaging of single DSBs in facultative heterochromatin (**Fig.2A**) using fluorescently tagged Mu2 to visualize DSBs, and fluorescently tagged polyhomeotic-proximal (ph-p) to visualize polycomb bodies (**Fig.2B**). Mu2 is the *Drosophila* ortholog of mammalian MDC1, and directly binds γH2Av^41,42^, while ph-p is one of the four core subunits of the *Drosophila* PRC1 (Polycomb Repressive Complex 1) and is enriched in fly polycomb bodies^43,44^ (**Fig.2B**). Strikingly, our live imaging analyses revealed that the majority of Mu2 foci (60%) that appear within polycomb bodies move outside the domain within ten minutes (**Fig.2C, D, Fig.S2A**), while euchromatic DSBs that appear outside polycomb bodies remain outside until their disappearance (**Fig.S2B**). Importantly, these movements are not specific for I-SceI-induced DSBs, since inducing DSBs in larval tissue using 5Gy gamma-radiation resulted in similar DSB dynamics in ph-p marked polycomb bodies (**Fig.2E, F**). The majority of radiation-induced Mu2 foci that appeared within polycomb bodies moved outside this domain within ten minutes after appearance, whereas euchromatic DSBs are resolved outside polycomb bodies (**Fig.S2C**). Taken together, our live imaging data demonstrate for the first time that DSBs move outside polycomb bodies to continue repair.

**Figure 2.**
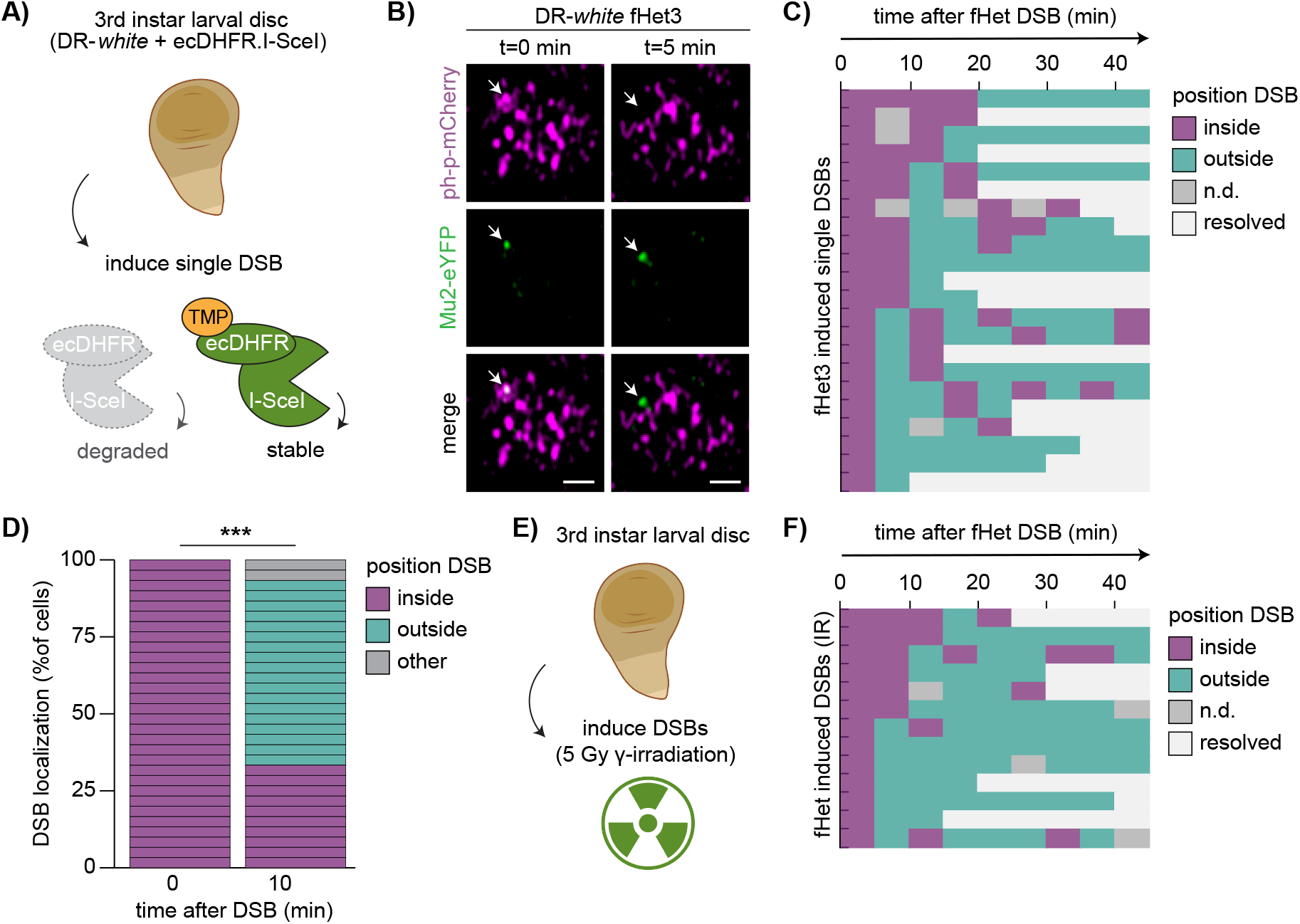
DSBs in facultative heterochromatin move outside polycomb bodies. **A**) Set up to analyze the dynamics of a single DSB in larval tissue. Wing discs of DR-*white*/ecDHFR-I-SceI 3^rd^ instar larvae were dissected and placed in medium with trimethoprim (TMP) to stabilize ecDHFR-I-SceI. **B**) Representative images of Mu2-eYFP focus dynamics (DSB marker, green) within the ph-p-mCherry domain (polycomb marker, magenta) in 3^rd^ instar larval wing disc. Arrow indicates Mu2-eYFP focus arising within ph-p-mCherry domain (0 min) and moving outside this domain (5 min). Scale bar = 1μm. **C**) Quantification of live imaging of Mu2-eYFP (DSB) dynamics and kinetics over time relative to the ph-p-mCherry domain using single DSB induction (ecDHFR-I-SceI, *fHet*3). Each row indicates one single Mu2-eYFP focus that appeared (0 min) within a polycomb body. Mu2-eYFP foci were followed up to 45 minutes after appearance. Colors indicate localization of the Mu2-eYFP focus (inside polycomb body [purple], outside [green], not detectable [grey], or resolved [white]). **D**) Quantification of Mu2-eYFP (DSB) localization 10 minutes after Mu2-eYFP appearance in ph-p domains (*fHet1* and *fHet3*). **E**) Set up to analyze the dynamics of DSBs using irradiation. Wing discs of 3^rd^ instar larvae were dissected and placed in medium, followed by exposure to 5Gy gamma-radiation (IR). **F**) Quantification of live imaging of Mu2-eYFP (DSB) dynamics and kinetics over time using 5Gy IR (quantification as in **C**). (***) p-value≤0.001, Chi-square test (**D**).

### Reduction in H3K27me3 at facultative heterochromatic DSBs is mediated by the histone demethylase dUtx

Considering the compact and silent state of facultative heterochromatin, we hypothesized that local chromatin changes could coincide with the specific DSB movements in polycomb bodies. To this end, we assessed the levels of the canonical facultative heterochromatin histone modification H3K27me3 by ChIP-qPCR at DSB sites (**Fig.1F, 3A**). We observed a decrease in H3K27me3 (loss of 22-34%) at two of the three heterochromatic DSB sites after I-SceI induction, while H3K27me3 at euchromatic DSB sites remained unchanged (**Fig.3A**). Since 16-18% of cells obtain a single DSB six hours after hsp.I-SceI induction (**Fig.S1B**), this decrease in H3K27me3 levels suggests a significant loss of H3K27me3 at individual heterochromatic DSBs. To exclude that the observed reduction in H3K27me3 is due to histone loss at the break site, we performed ChIP-qPCR for histone H3, which did not reveal any significant differences in histone H3 levels at euchromatic and heterochromatic DSB sites (**Fig.3B, Fig.S3A**). The reduction in H3K27me3 was observed in two of the three heterochromatic integrations, suggesting that not all heterochromatic DSBs induce evident loss of H3K27me3. A possibility is that our ChIP assay may not be sensitive enough to pick up on subtle differences in H3K27me3 levels. Nevertheless, our data reveal that DSBs in facultative heterochromatin are frequently accompanied by a local reduction in H3K27me3.

**Figure 3.**
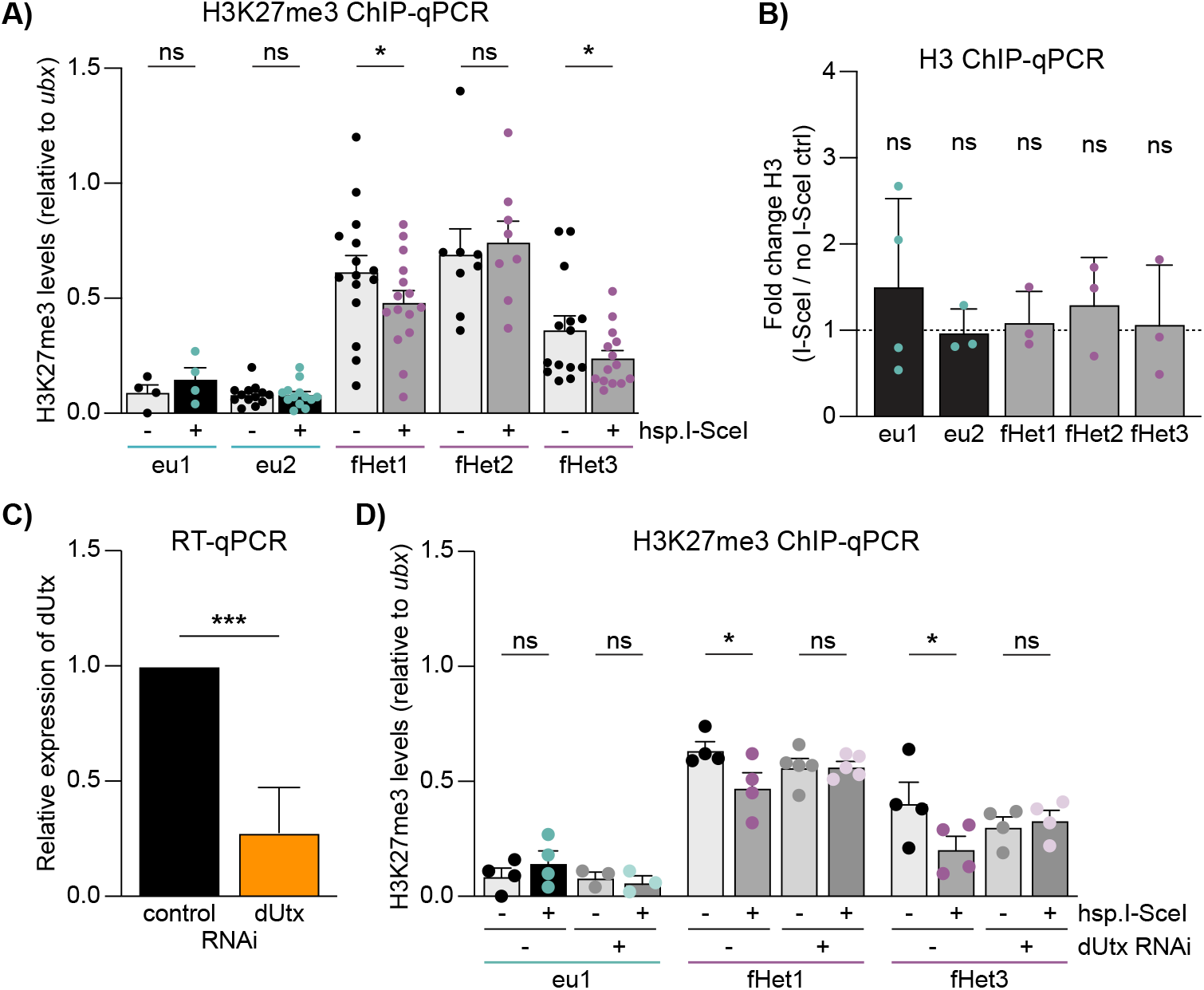
Loss of H3K27me3 at facultative heterochromatic DSBs is mediated by dUtx. **A**) ChIP-qPCR analysis for H3K27me3 in the absence (-) and presence (+) of hsp.I-SceI (as in Fig.1F) at indicated DR-*white* integration sites. H3K27me3 levels were normalized using a *ubx* qPCR primer set as an internal positive control (*ubx* has consistently high H3K27me3 levels). Averages are shown for ≥4 independent experiments +SEM. **B**) ChIP-qPCR analysis for histone H3. Fold change indicates H3 levels at DR-*white* loci in hsp.I-SceI-expressing larvae (+DSB) divided by larvae not expressing hsp.I-SceI (-DSB). Dooed line indicates no change (fold change = 1). H3 levels were normalized using a *yellow* qPCR primer as an internal positive control. Bars indicate fold change +SD of ≥3 independent experiments. **C**) Relative dUtx expression level in 3^rd^ instar larvae determined using RT-qPCR and normalized to an internal control gene (tubulin). Bars indicate averages +SD of 8 single larvae per condition (luciferase control RNAi or dUtx RNAi). **D**) ChIP-qPCR analysis for H3K27me3 in the absence or presence of a DSB (-/+ hsp.I-SceI), with or without dUtx RNAi depletion (as in Fig.1F). H3K27me3 levels were normalized using a *ubx* qPCR primer set as internal positive control. Averages are shown for ≥4 independent experiments +SEM. (ns) not significant, (*) p-value≤0.05, (***) p-value ≤0.001, paired t-test (**A-D**).

We hypothesized that the reduction in H3K27me3 levels at heterochromatic DSBs could be mediated by a histone demethylase that actively removes the methyl groups from H3K27. In *Drosophila*, dUtx is the only protein described to demethylate H3K27me3^45^. To determine whether dUtx removes the methyl group at heterochromatic DSB, we depleted dUtx using RNAi in 3^rd^ instar larvae (**Fig.3C**) and assessed H3K27me3 levels at the DSB sites using ChIP-qPCR. Indeed, dUtx depletion leads to retention of H3K27me3 at DSBs in heterochromatin, whereas the H3K27me3 levels at euchromatic DSBs remain unaffected upon DSB induction in the presence or absence of dUtx (**Fig.3D**). Loss of dUtx did not alter the levels of the DSB marker γH2Av, indicating that the retention of H3K27me3 at DR-*white* sites upon dUtx depletion is not due to inefficient cutting by I-SceI (**Fig.S3B**). These data suggest that dUtx mediates the removal of H3K27me3 at DSB sites specifically in facultative heterochromatin.

### DSB movement and HR repair in facultative heterochromatin depend on dUtx

We next hypothesized that the local loss of H3K27me3 at heterochromatic DSBs could be required to promote DSB movement outside polycomb bodies. To test this, we turned to *Drosophila* cells in culture which allow for in-depth visualization of repair processes in combination with RNAi-mediated depletions. We visualized the dynamics of the early HR protein ATR Interacting Protein (ATRIP) within polycomb bodies (ph-p domains) in the presence or absence of dUtx using live imaging of 5Gy gamma-radiated *Drosophila* Kc cells (**Fig.4A**). ATRIP binds to RPA-coated ssDNA overhangs, which are produced early in HR during 5’ to 3’ end resection of the DSB^46^. We find that upon irradiation of control cells, ATRIP foci appear inside polycomb bodies and move outside these domains within ten minutes (**Fig.4B, C, Fig.S4A**), which recapitulates our findings on Mu2 foci dynamics at heterochromatic DSBs in larval tissues (**Fig.2C, D, F**). Interestingly, ATRIP gets recruited inside polycomb bodies upon irradiation in the absence of dUtx, suggesting that end resection steps are independent of dUtx (**Fig.4D, E, Fig.S4B**). However, dUtx depletion does lead to a delay in the movement of ATRIP-coated DSBs outside polycomb bodies. In control cells, 77% of ATRIP foci move outside the polycomb bodies within 10 minutes, while in dUtx depleted cells only 53% move out within this timeframe (**Fig.4C**). In line with this, we find an increased accumulation of ATRIP foci within the ph-p domains (21% in control cells during the course of our imaging experiment, compared to 33% in dUtx-depleted cells) indicative of defects in DSB movement upon dUtx depletion (**Fig.4E**). Together, these results suggest that the early steps of HR (end resection, ATRIP recruitment) occur efficiently within polycomb bodies and are independent of H3K27me3 demethylation at the DSB site. However, dUtx-mediated demethylation of H3K27me3 is required for the subsequent DSB movement outside the polycomb body.

**Figure 4.**
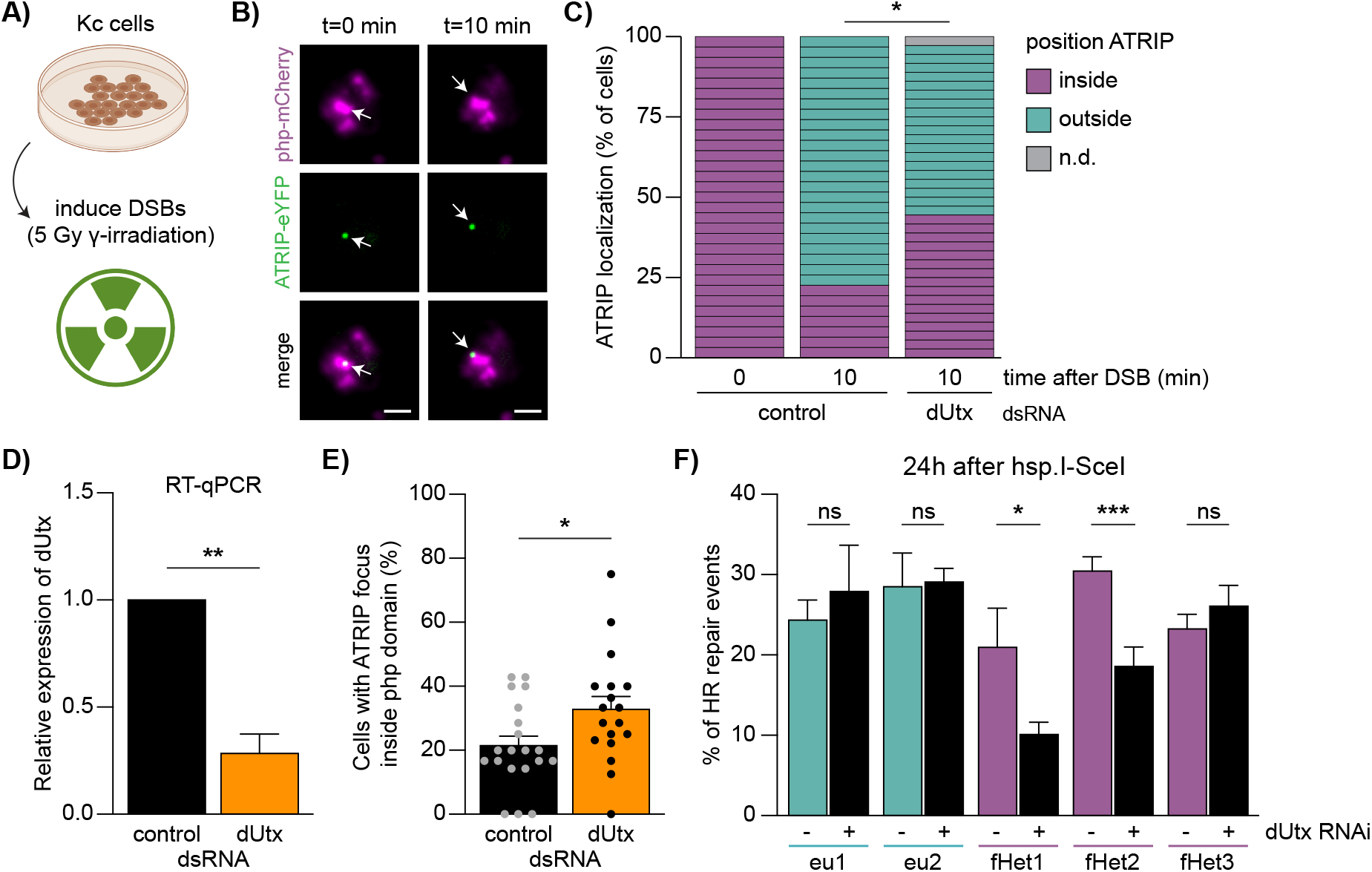
DSB movement and HR repair in facultative heterochromatin depend on dUtx. **A)** Cell culture set up to analyze the dynamics of ATRIP localization upon DSB induction using 5Gy gamma-radiation. **B**) Representative images of ATRIP-eYFP focus (green) dynamics within the ph-p-mCherry domain (magenta). Arrow indicates ATRIP-eYFP focus arising within ph-p-mCherry domain (0 min) and moving outside this domain (10 min). Scale bar = 2μm. **C**) Quantification of ATRIP-eYFP (DSB) localization 10 minutes after ATRIP-eYFP appearance, ≥3 independent experiments per condition (yellow control dsRNA or dUtx dsRNA). **D**) Relative dUtx expression level in *Drosophila* Kc cells normalized to an internal control gene (tubulin), determined using RT-qPCR. Bars indicate averages +SD of 3 independent experiments per condition. **E**) Quantification of the number of cells with an ATRIP-eYFP focus inside ph-p-mCherry domain divided by the total number of cells expressing both ATRIP-eYFP and ph-p-mCherry. Each dot indicates the percentage of cells with ATRIP foci in ph-p per acquired image (3 independent experiments with ≥5 acquired images per condition). Bars indicate averages +SEM. **F**) DR-*white*/hsp.I-SceI larvae with (+) or without (-) dUtx RNAi were heat-shocked for one hour to induce I-SceI. Repair PCR products were Sanger sequenced 24 hours after I-SceI induction and analyzed using the TIDE algorithm. Graph shows percentage of identified HR products. Bars indicate averages +SEM of ≥5 independent experiments (larvae) per condition. (ns) not significant, (*) p-value≤0.05, (**) p-value≤0.01, (***) p-value≤0.001, Chi-square test (C), paired t-test (D) and unpaired t-test (**E, F**).

To test whether this dUtx-dependent DSB movement is required for later repair steps in facultative heterochromatin, we employed our *in vivo* reporter system, which allows the direct assessment of DSB repair pathway choice (HR/NHEJ) by sequencing repair products (**Fig.1A**). Strikingly, loss of dUtx in larvae revealed a 39-52% relative reduction in the proportion of HR repair products at two of the three heterochromatic DSB sites, while euchromatic DSB repair products remained unchanged (**Fig.4F**). This reduction in HR repair was accompanied by an increase in NHEJ repair (**Fig.S4C**). These changes in DSB repair pathway choice are not due to indirect cell-cycle effects, since dUtx depletion did not significantly affect cell cycle progression in the Fly-FUCCI system^47^ (**Fig.S4D, E**).

Although dUtx depletion did not affect relative HR levels at the *fHet3* integration (**Fig.4F**), we do find that loss of dUtx results in a reduction in the total number of identified repair products at the *fHet3* integration, as well as at the *fHet2* integration site (**Fig.S4F**). A reduction in identified repair products is indicative of defects or delays in DSB repair. Interestingly, this reduction in total identified repair products is more evident when strictly assessing repair of the *fHet3* region in wing disc tissues, which have strong silencing of genes nearby the *fHet3* integration site (e.g. hmx)^48^. This indicates that DSB sites with high H3K27me3 levels depend more heavily on dUtx for repair (**Fig.S4G**). Indeed, brain tissues with high gene expression levels (low H3K27me3) nearby *fHet3* do not show this reduction in repair efficiency upon dUtx depletion. Together, these results suggest that DSB repair regulation is defective at all heterochromatic sites in the absence of dUtx.

Finally, although DSBs at the *fHet2* locus did not induce H3K27me3 loss (**Fig.3A**) this region showed a clear defect in HR in the absence of dUtx (**Fig.4F**). This suggests that dUtx plays an important role in DSB repair at this integration site and we may not have been able to identify dUtx-mediated loss of H3K27me3 levels at this site due to compensatory histone methylation activities or relatively low sensitivity of our ChIP assay. Altogether, our results reveal that early HR steps in facultative heterochromatin can be performed in the absence of dUtx, and that dUtx is specifically required for DSB movement and completion of repair.

To assess the physiological role of dUtx in DSB repair in facultative heterochromatin, we wished to determine whether development of flies mutant for dUtx depends on the presence of intact DNA damage signaling. To do so, we crossed heterozygous dUtx mutant flies with flies that contain a truncation mutation in Ataxia telangiectasia and Rad3 related (ATR, mei41). ATR is one of the earliest kinases that rapidly respond to DSB events^49^. Combining dUtx mutant flies with an ATR mutation reduces relative viability by 16%, indeed suggesting that dUtx mutant flies depend on correct DNA damage repair signaling for their development (**Fig.S4H**). This result reveals the physiological role of dUtx and suggests that in the absence of proper facultative heterochromatic DSB repair, flies depend on active DNA damage checkpoint signaling to maintain viability.

## DISCUSSION

Chromatin forms dynamic domains in the nucleus, each characterized by specific molecular properties, which can directly influence the DSB response. However, how transcriptionally inactive facultative heterochromatin (i.e. polycomb chromatin) influences DSB repair remains poorly understood. To address this question and understand how eukaryotic cells maintain the integrity of silenced developmental genes, we here integrated inducible single DSB systems in euchromatin and facultative heterochromatin in *Drosophila Melanogaster*. This allowed us to comprehensively study DSB repair in facultative heterochromatin in animal tissue for the first time. We find that DSBs in facultative heterochromatin rapidly move outside polycomb bodies within minutes after their appearance. This movement depends on the H3K27me3 demethylase dUtx. In line with this, we find evidence for dUtx-mediated loss of the silencing mark H3K27me3 near DSBs in facultative heterochromatin. Our data further reveal that early steps of HR (i.e. end resection) can occur efficiently within polycomb bodies and are independent of dUtx, whereas dUtx is required to promote subsequent DSB movement and the completion of HR. Together, we propose a model in which resected DSBs in polycomb bodies are subjected to dUtx-mediated loss of the silencing mark H3K27me3, which in turn promotes DSB movement and timely repair by homologous recombination (**Fig.5**).

**Figure 5.**
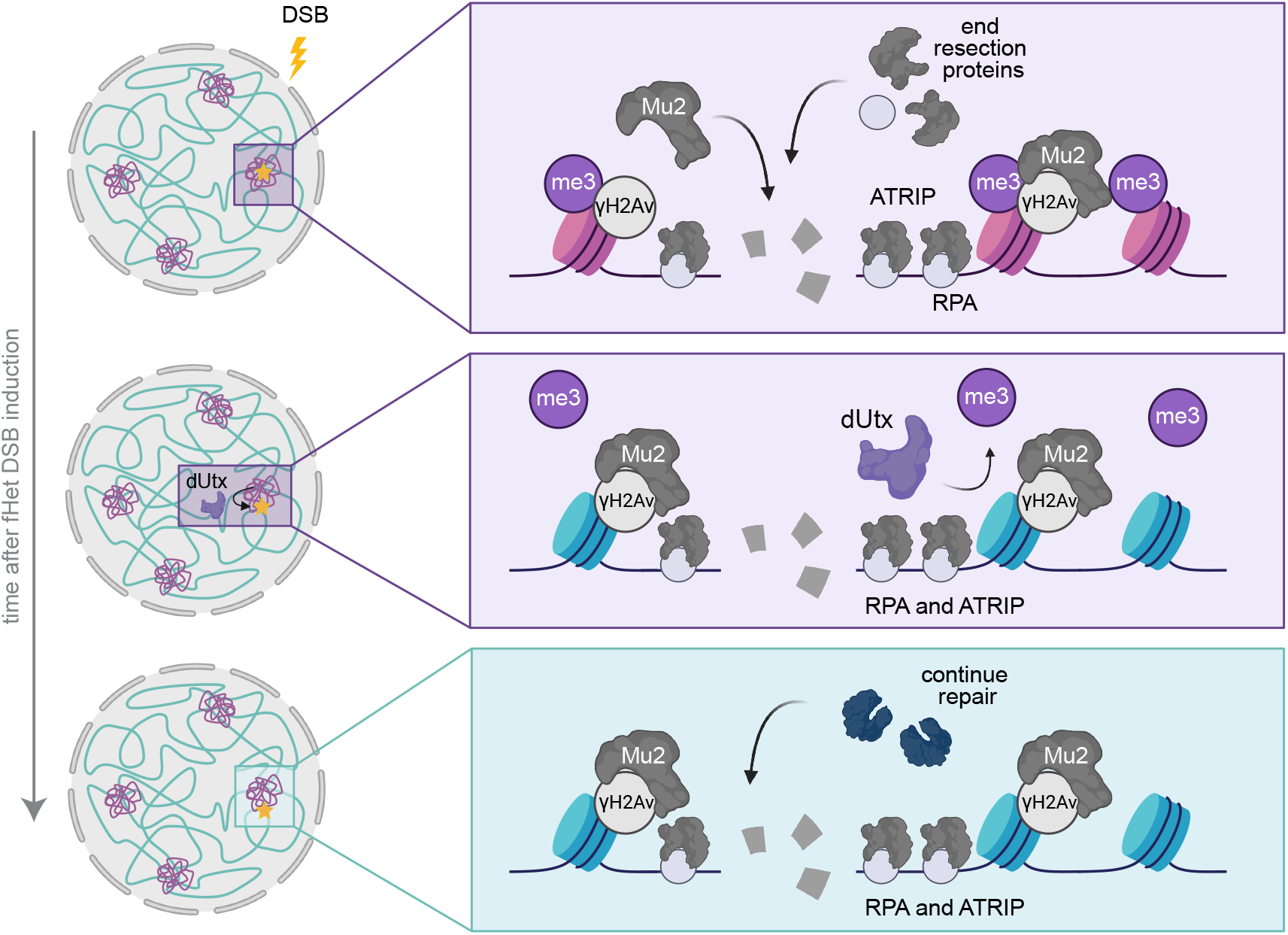
Model. Model for facultative heterochromatin DSB repair. Facultative heterochromatin (purple) and euchromatin (blue) form distinct domains within the nucleus. Facultative heterochromatin is characterized by H3K27me3 and accumulates in polycomb bodies. DSBs in facultative heterochromatin undergo early steps of HR (i.e. Mu2-recruitment, end resection) within polycomb bodies. dUtx is required to demethylate H3K27me3 at the DSB site and promote the subsequent DSB movement to efficiently resolve DSB using HR.

Specific movements of DSBs have been identified to occur in a variety of chromatin compartments including centromeres^16^, nucleoli^17–19^ and pericentromeric constitutive heterochromatin^16,20–22^. Movement of these DSBs has been suggested to promote binding of HR proteins^17,18^, as well as prevent aberrant recombination between repetitive sequences^20,21^. Here, we find for the first time that DSBs move outside polycomb bodies *in vivo* (**Fig.2**). In contrast to centromeres, nucleoli and constitutive heterochromatin, facultative heterochromatin mainly contains unique sequences and is deprived of repetitive sequences. We therefore propose that DSB movement in facultative heterochromatin did not evolve to prevent aberrant recombination, but rather reflects the necessity to create a repair-competent state, facilitating access to the DSB repair machinery (**Fig.5**).

Our data suggest that the movement of DSBs outside polycomb bodies is directly regulated by the dUtx-mediated removal of H3K27me3 at the break site (**Fig.3, 4**). H3K27me3 is required to recruit PcG proteins to enhance compaction and maintain a silenced state^9–13^. Therefore, active removal of H3K27me3 at the DSB site by dUtx could directly lead to a local loss of PcG proteins. This can subsequently lead to changes in the molecular- and biophysical-properties of the DSB locus, creating an environment distinct from the surrounding polycomb body and the active expulsion or passive separation of the DSB from the polycomb body. In line with this hypothesis, we find that in the absence of dUtx, and the subsequent retention of H3K27me3, DSBs remain longer within the polycomb body (**Fig.4**). These results reveal analogies with our previous findings at DSBs in constitutive heterochromatin, where we found that loss of the silencing mark H3K9me3 at DSBs by the histone demethylase dKDM4A ensures DSB movement outside the constitutive heterochromatin domain^32,33^.

Since we did not observe a complete inhibition of facultative heterochromatic DSB movement upon dUtx depletion, additional processes are likely involved, such as DSB end processing or the recruitment of specific chromatin-or repair-proteins. End resection as well as chromatin proteins, including the cohesin- and SMC5/6-complexes, drive DSB movement in other chromatin domains^16,18,20^, suggesting that additional components could be driving movements in polycomb bodies.

Our findings indicate that dUtx-mediated demethylation of H3K27me3 at facultative heterochromatic DSBs is important for repair pathway choice, since dUtx loss shiis the choice towards NHEJ, resulting in decreased HR (**Fig.4**). Considering that dUtx depletion only affects DSB repair pathway choice in facultative heterochromatin, not euchromatin, these repair pathway changes are unlikely to be driven by indirect general defects in cell cycle progression (**Fig.S4**) or transcriptional regulation of repair genes^50^. Therefore, we hypothesize that the HR/NHEJ repair pathway choice at facultative heterochromatin could be directly regulated by dUtx through two non-mutually exclusive mechanisms: (1) defects in DSB movement, and (2) direct impact on binding of HR-or NHEJ-proteins at DSB sites.

In the first model, dUtx is promoting HR by moving the DSB to a more HR-prone chromatin state depleted of silencing marks. The movement might therefore specifically facilitate the access to ‘late’ HR repair proteins (e.g. Rad51 loading or helicases to resolve D-loops), usually excluded from the compact polycomb state. Moreover, moving an HR-proficient DSB away from the compact facultative heterochromatin might provide the required chromatin mobility necessary to perform homology search^51^. Indeed, we observe that loss of dUtx has no impact on the initial stages of HR, such as end resection and ATRIP loading, in polycomb bodies. However, dUtx loss does impede DSB movement, and subsequent later HR steps as evidenced by the decreased number of HR repair products identified at facultative heterochromatic DSB sites (**Fig.4**).

In our second model, we propose that the decreased frequency of HR repair in the absence of dUtx is caused by changes in binding of repair proteins to histone modifications at the break site. It is possible that H3K27me3 is able to directly bind specific NHEJ proteins within the polycomb body. Alternatively, H3K7me1 or unmethylated H3K27 residues, as a result of dUtx-mediated H3K27me3 demethylation, could directly recruit proteins important for HR. In line with this hypothesis, previous work identified that the HR-promoting TONSL-MMS2L complex has a higher affinity for unmodified histones, generated following DNA replication^52^.

Despite differences in dUtx-dependency for repair pathway choice, we find the frequencies of HR and NHEJ repair pathway usage in facultative heterochromatin and euchromatin to be similar in wild type *Drosophila* (**Fig.1, Fig.S1**). These results are consistent with previous findings in which DSBs in H3K27me3-enriched imprinted loci in mice did not differ in repair pathway usage when compared to the corresponding active allele^53^. In addition, both HR and NHEJ components are recruited to laser-damaged inactive X chromosomes in female human cells^25^. In contrast, a recent study that used a sequencing-based reporter system in cancer cells to investigate the impact of chromatin on CRISPR-Cas9-induced DSBs did reveal differences in repair pathway usage in H3K27me3-enriched regions^54^. The authors found a relative decrease in NHEJ and concurrently an increase in usage of Microhomology-Mediated End-Joining (MMEJ) within H3K27me3-enriched regions. MMEJ is an error-prone mechanism that relies on short-range end resection and uses homologous sequences to align the broken ends. In contrast to our results, these findings suggest that end resection-based repair pathways are preferred at facultative heterochromatic DSBs. These different outcomes could indicate differences in repair pathway usage between species or could be due to differences in the approach used to induce DSB induction (CRISPR-Cas9 versus I-SceI). Moreover, heterochromatin properties *in vivo* may vary from that observed in cultured tumor cells, potentially leading to disparate outcomes.

In conclusion, our work demonstrates that DSBs in facultative heterochromatin require specific local chromatin changes and DSB movements for their faithful repair in animal tissue. Our results emphasize the importance of understanding how different chromatin components influence DSB repair pathway choice and maintain genome stability across diverse chromatin domains. Facultative heterochromatin regions are often associated with high mutational loads in cancer^55^, indicating that these domains are particularly vulnerable to aberrant DNA damage repair. Moreover, the human homolog of dUtx, UTX, is often mutated in cancer^56^. In the long-term, research into DNA damage repair in heterochromatin will give insights into how misregulation of chromatin proteins, such as UTX, could result in increased genome instability and specific mutational signatures in cancer, ultimately contributing to disease development.

## METHODS

### Constructs

The DR-*white* construct was created previously^22^. ph-p was N-terminally tagged with mCherry and cloned in a pCasper5 vector for random p-element transformation in flies. For cell culture experiments, ph-p and ATRIP were cloned into pCopia vectors containing N-terminal mCherry or C-terminal eYFP epitope tags respectively. ph-p was cloned from the pFastBac plasmid (Addgene #1925), whereas ATRIP was cloned from cDNA generated from RNA extracted from wild type flies.

### Fly lines

Flies were grown at room temperature on standard medium, except otherwise specified. Embryo injection and generation of new DR-*white* and ph-p-mCherry fly lines were performed by BestGene, Inc. DR-*white* aoB containing plasmids were integrated in Minos-mediated integration casseoe (MiMIC) integration sites as described previously^22^. Facultative heterochromatin integration sites were selected based on high H3K27me3 levels in OregonR flies (modEncode) and near a gene known to be regulated by H3K27me3 and/or PcG proteins^57^. Euchromatin integration sites were selected based on low H3K27me3 and low H3K9me2/3 levels. To create ph-p-mCherry fly lines, pCasper5-ph-p-mCherry plasmid containing the copia promoter and P-element transposons was injected in embryos of w1118 flies by Bestgene (Chino Hills, CA, USA). To induce knockdown of either CtIP or dUtx, flies containing Gal4 driven by an Actin5C promoter were crossed with UAS-CtIP RNAi or UAS-dUtx RNAi. A list of MiMIC integration sites to generate the DR-*white* fly lines as well as all fly lines used can be found in Supplemental Table S1.

### ChIP-qPCR

Third instar DR-*white* larvae (+ and -hsp.I-SceI) were heat-shocked for 1 hour to express hsp.I-SceI. Six hours after I-SceI activation, 40 larvae per condition were pooled together to extract chromatin following the ChIP protocol of the Kevin White lab (https://www.encodeproject.org/documents/f890fde6-924c-4265-a60f-c5810401066d/), with slight adjustments. In short, larvae were homogenized in buffer A1 (60mM KCl, 15mM NaCl, 15mM HEPES pH7.6, 4mM MgCl2, 0.5% TritonX-100, 0.5mM DTT, protease inhibitor (Roche #1873580)) and fixed using 1.8% paraformaldehyde (EMS). Fixation was stopped by addition of glycine. After several washing steps with buffer A1, nuclei were isolated (140mM NaCl, 15mM HEPES pH7.6, 1mM EDTA, 0.5mM EGTA, 0.1% sodium deoxycholate, 1% TritonX-100, 500μM DTT, protease inhibitors, 0.1% SDS and 0.5% N-lauroylsarcosine) and the chromatin was fragmented by sonication for 10 cycles on Bioruptor (Diagenode high settings, 30’’ on/off). Chromatin was separated from cell debris using centrifugation and stored at -80C for a maximum of 2 months.

ChIP was performed as described earlier^58^ using 1-2 ug chromatin and 1-5 ug antibody. ChIP antibodies used were mouse anti-γH2Av (Developmental Studies Hybridoma Bank, UNC93-5.2.1), rabbit anti-H3K27me3 (Invitrogen, MA5-11198), and rabbit anti-H3 (Abcam, ab1791). Enrichment of the specific histone mark was quantified by qPCR using SYBR Green Master Mix (Roche) and qPCR primers 1.4 kb away from the DSB (3xp3) as well as primers for an internal control. All qPCR primer sequences are listed in Supplemental Table S1.

Epicypher SNAP-ChIP K-MetStat panel (19-1001) was used to validate the specificity of the H3K27me3 antibody used for ChIP (**Fig.S3C**). The SNAP-ChIP panel contains barcoded nucleosomes with a specific methylation mark (unmethylated H3, H3K4me1/me2/me3, H3K9me1/me2/me3, H3K27me1/me2/me3, and H3K36me1/me2/me3). 0.4μL of 0.6 nM K-MetStat stock per 2 ug of chromatin was added at the start of the ChIP procedure. Subsequent qPCR analysis using primers specific to unique barcodes corresponding to each modified nucleosome in the panel allows the quantification of antibody specificity and efficiency.

### DR-white repair product analysis

Quantification of somatic repair products in DR-*white*, I-SceI larvae was performed as described previously^22^. In short, either hsp.I-SceI was expressed using heat-shock in third instar DR-*white*/hsp.I-SceI larvae, 24 hours before single larvae were collected. Alternatively, ecDHFR-I-SceI was expressed by feeding DR-*white*/ecDHFR-I-SceI 80μM trimethoprim (Sigma) throughout development (3-4 days), thereby stabilizing the ecDHFR-I-SceI protein. To prepare trimethoprim containing food, 1.67g instant *Drosophila* medium (Formula 4-24, Carolina Biological Supply) was mixed with 5 mL non distilled water containing 5.3 uL of 100mM trimethoprim while vortexing.

To analyze repair products, the upstream *white* gene was PCR amplified and the PCR product was treated with ExoSAP-IT to enzymatically remove excess primers and unincorporated nucleotides, followed by Sanger sequencing. Analysis of Sanger sequences was performed using the TIDE (tracking of indels by decomposition) algorithm^36^. HR repair products were identified by loss of the I-SceI cleavage site and appearance of the wildtype *white* gene, which is essentially a 23-nucleotide deletion at the I-SceI cut site. NHEJ products were identified as insertions and deletions up to 25bp, except for the 23-nucleotide deletion. For PCR and sequencing primers, see Supplemental Table S1.

### Immunofluorescence staining

Wing discs were dissected from third instar DR-*white* larvae and fixed on slides as described earlier^59^ . Slides were stored in 96% ethanol at -20°C until staining procedure. Slides were thawed at room temperature and washed in PBS for 20 minutes. Tissues were blocked using 0.4% Triton in PBS and 5% milk for 1 hour at room temperature. Primary antibody incubations were performed overnight at 4°C in block buffer. Primary antibody used for imaging was mouse anti-γH2Av (1:250, Developmental Studies Hybridoma Bank, UNC93-5.2.1). Slides were washed 3 times with block buffer. Secondary antibody incubation was performed at room temperature in PBS 0.4% Triton for 2 hours. Secondary antibody used was Alexa 568 goat anti-mouse (1:600; Invitrogen). Slides were subsequently washed 3 times with 0.4% Triton in PBS, incubated with 3μg/ml DAPI for 30 minutes, washed with PBS and mounted using Prolong Diamond Antifade Mountant and a 20x20 mm #1.5 coverslip.

### Cell culture

Kc cells were cultured in CCM3 medium (Avantor) supplemented with Antibiotic Antimycotic Solution (Sigma) at 27°C. To induce expression of ph-p-mCherry and ATRIP-eYFP, Kc cells were transiently transfected with 400ng of each plasmid using the TransIT-2020 reagent (Mirus). Live imaging was performed 72 hours after transfection. To induce RNAi-mediated depletion of dUtx, cells were transfected with 5ug dsRNA (TransIT-2020 reagent (Mirus)) and harvested or subjected to imaging 3 days later. dsRNA was generated using a MEGAScript T7 transcription kit (Life Technologies). PCR products containing a T7 promoter sequence and the target regions were used as templates (Supplemental Table S1).

### RT-qPCR

RNA was isolated by homogenizing either single larvae or Kc cells in 200μL Trizol (Invitrogen) using an electrical douncer (VWR). After addition of 40μL of chloroform and centrifugation, RNA from the aqueous phase was precipitated using isopropanol and further purified using an ethanol wash step. cDNA was synthesized using iScript following standard cDNA synthesis protocol (Bio-Rad). qPCR was subsequently performed on cDNA with gene-specific primers (Supplemental Table S1).

### Imaging

Images of fixed tissue, Fly-FUCCI wing discs and transfected Kc cells were acquired using a 60x oil immersion objective (NA 1.42) on a DeltaVision microscope (DeltaVision Spectris; Applied Precision, LLC). For all DSB tracking experiments in wing discs, time-lapse images were acquired using a LD C-Apochromat 63x/1.15 W Korr M27 objective on a LSM880 microscope with Airyscan (Zeiss), and images were processed using the Zeiss ZEN software. Time-lapse images were acquired once every 5-10 minutes. Image analysis and focus tracking were performed manually using the Fiji image analysis software.

For live imaging of wing discs, third instar larvae were dissected and wing discs were placed on a slide in 10 μL of Schneider S1 medium supplemented with 10% FBS and covered with a 20×20 mm #1.5 coverslip as described previously^60^. To induce ecDHFR-I-SceI in larval tissue, 400 μM trimethoprim (Sigma) was added 15 minutes prior to imaging.

## ACKNOWLEDGEMENTS

We thank Jurian Schuijers and Lucie van Leeuwen for critically reading our manuscript and all the members of the Lens and Janssen laboratories for their valuable input during lab meetings. Special thanks to Johannes Lehmann for his advice on live imaging analysis. This work was funded by the European Research Council (ERC) under the European Union’s Horizon 2020 research and innovation program, grant agreement No. 850405, and VIDI VI.Vidi.203.001 financed by the Dutch Research Council (NWO). Several figures were created with BioRender.com.

## REFERENCES

1. Janssen, A. & Medema, R. H. Genetic instability: tipping the balance. Oncogene 32, 4459–4470 (2013).

2. Scully, R., Panday, A., Elango, R. & Willis, N. A. DNA double-strand break repair-pathway choice in somatic mammalian cells. Nat. Rev. Mol. Cell Biol. 20, 698–714 (2019).

3. Clouaire, T. & Legube, G. A Snapshot on the Cis Chromatin Response to DNA Double-Strand Breaks. Trends Genet. 35, 330–345 (2019).

4. van Bueren, M. A. E. & Janssen, A. The impact of chromatin on double-strand break repair: Imaging tools and discoveries. DNA Repair (Amst) 133, 103592 (2023).

5. Lewis, E. B. A gene complex controlling segmentation in Drosophila. Nature 276, 565–570 (1978).

6. Trojer, P. et al. L3MBTL1, a histone-methylation-dependent chromatin lock. Cell 129, 915–928 (2007).

7. Messmer, S., Franke, A. & Paro, R. Analysis of the functional role of the Polycomb chromo domain in Drosophila melanogaster. Genes Dev. 6, 1241–1254 (1992).

8. Buchenau, P., Hodgson, J., Strutt, H. & Arndt-Jovin, D. J. The distribution of polycomb-group proteins during cell division and development in Drosophila embryos: impact on models for silencing. J. Cell Biol. 141, 469–481 (1998).

9. Wani, A. H. et al. Chromatin topology is coupled to Polycomb group protein subnuclear organization. Nat. Commun. 7, 10291 (2016).

10. Grimaud, C. et al. RNAi components are required for nuclear clustering of Polycomb group response elements. Cell 124, 957–971 (2006).

11. Lanzuolo, C., Roure, V., Dekker, J., Bantignies, F. & Orlando, V. Polycomb response elements mediate the formation of chromosome higher-order structures in the bithorax complex. Nat. Cell Biol. 9, 1167–1174 (2007).

12. Bantignies, F. et al. Polycomb-dependent regulatory contacts between distant Hox loci in Drosophila. Cell 144, 214–226 (2011).

13. Isono, K. et al. SAM domain polymerization links subnuclear clustering of PRC1 to gene silencing. Dev. Cell 26, 565–577 (2013).

14. Aymard, F. et al. Transcriptionally active chromatin recruits homologous recombination at DNA double-strand breaks. Nat. Struct. Mol. Biol. 21, 366–374 (2014).

15. Aymard, F. et al. Genome-wide mapping of long-range contacts unveils clustering of DNA double-strand breaks at damaged active genes. Nat. Struct. Mol. Biol. 24, 353–361 (2017).

16. Tsouroula, K. et al. Temporal and Spatial Uncoupling of DNA Double Strand Break Repair Pathways within Mammalian Heterochromatin. Mol. Cell 63, 293–305 (2016).

17. van Sluis, M. & McStay, B. A localized nucleolar DNA damage response facilitates recruitment of the homology-directed repair machinery independent of cell cycle stage. Genes Dev. 29, 1151–1163 (2015).

18. Marnef, A. et al. A cohesin/HUSH- and LINC-dependent pathway controls ribosomal DNA double-strand break repair. Genes Dev. 33, 1175–1190 (2019).

19. Torres-Rosell, J. et al. The Smc5-Smc6 complex and SUMO modification of Rad52 regulates recombinational repair at the ribosomal gene locus. Nat. Cell Biol. 9, 923–931 (2007).

20. Chiolo, I. et al. Double-strand breaks in heterochromatin move outside of a dynamic HP1a domain to complete recombinational repair. Cell 144, 732–744 (2011).

21. Ryu, T. et al. Heterochromatic breaks move to the nuclear periphery to continue recombinational repair. Nat. Cell Biol. 17, 1401–1411 (2015).

22. Janssen, A. et al. A single double-strand break system reveals repair dynamics and mechanisms in heterochromatin and euchromatin. Genes Dev. 30, 1645–1657 (2016).

23. Müller, I. et al. Species conserved DNA damage response at the inactive human X chromosome. Mutat. Res. 756, 30–36 (2013).

24. Żylicz, J. J. & Heard, E. Molecular Mechanisms of Facultative Heterochromatin Formation: An X-Chromosome Perspective. Annu. Rev. Biochem. 89, 255–282 (2020).

25. Chansard, A., Pobega, E., Caron, P. & Polo, S. E. Imaging the response to DNA damage in heterochromatin domains. Front. Cell Dev. Biol. 10, 920267 (2022).

26. Clouaire, T. et al. Comprehensive Mapping of Histone Modifications at DNA Double-Strand Breaks Deciphers Repair Pathway Chromatin Signatures. Mol. Cell 72, 250–262.e6 (2018).

27. Chou, D. M. et al. A chromatin localization screen reveals poly (ADP ribose)-regulated recruitment of the repressive polycomb and NuRD complexes to sites of DNA damage. Proc Natl Acad Sci USA 107, 18475–18480 (2010).

28. Abu-Zhayia, E. R., Awwad, S. W., Ben-Oz, B. M., Khoury-Haddad, H. & Ayoub, N. CDYL1 fosters double-strand break-induced transcription silencing and promotes homology-directed repair. J. Mol. Cell Biol. 10, 341–357 (2018).

29. Ayrapetov, M. K., Gursoy-Yuzugullu, O., Xu, C., Xu, Y. & Price, B. D. DNA double-strand breaks promote methylation of histone H3 on lysine 9 and transient formation of repressive chromatin. Proc Natl Acad Sci USA 111, 9169–9174 (2014).

30. Khurana, S. et al. A macrohistone variant links dynamic chromatin compaction to BRCA1-dependent genome maintenance. Cell Rep. 8, 1049–1062 (2014).

31. Rath, B. H., Waung, I., Camphausen, K. & Tofilon, P. J. Inhibition of the histone H3K27 demethylase UTX enhances tumor cell radiosensitivity. Mol. Cancer Ther. 17, 1070–1078 (2018).

32. Colmenares, S. U. et al. Drosophila Histone Demethylase KDM4A Has Enzymatic and Non-enzymatic Roles in Controlling Heterochromatin Integrity. Dev. Cell 42, 156–169.e5 (2017).

33. Janssen, A., Colmenares, S. U., Lee, T. & Karpen, G. H. Timely double-strand break repair and pathway choice in pericentromeric heterochromatin depend on the histone demethylase dKDM4A. Genes Dev. 33, 103–115 (2019).

34. Do, A. T., Brooks, J. T., Le Neveu, M. K. & LaRocque, J. R. Double-strand break repair assays determine pathway choice and structure of gene conversion events in Drosophila melanogaster. G3 (Bethesda) 4, 425–432 (2014).

35. Rogakou, E. P., Pilch, D. R., Orr, A. H., Ivanova, V. S. & Bonner, W. M. DNA doublestranded breaks induce histone H2AX phosphorylation on serine 139. J. Biol. Chem. 273, 5858–5868 (1998).

36. Brinkman, E. K., Chen, T., Amendola, M. & van Steensel, B. Easy quantitative assessment of genome editing by sequence trace decomposition. Nucleic Acids Res. 42, e168 (2014).

37. Plys, A. J. et al. Phase separation of Polycomb-repressive complex 1 is governed by a charged disordered region of CBX2. Genes Dev. 33, 799–813 (2019).

38. Cheutin, T. & Cavalli, G. Progressive polycomb assembly on H3K27me3 compartments generates polycomb bodies with developmentally regulated motion. PLoS Genet. 8, e1002465 (2012).

39. Tatavosian, R. et al. Nuclear condensates of the Polycomb protein chromobox 2 (CBX2) assemble through phase separation. J. Biol. Chem. 294, 1451–1463 (2019).

40. Sun, W. et al. Histone marks enable formation of immiscible phase-separated chromatin compartments. BioRxiv (2023) doi:10.1101/2023.12.02.569699.

41. Stucki, M. et al. MDC1 directly binds phosphorylated histone H2AX to regulate cellular responses to DNA double-strand breaks. Cell 123, 1213–1226 (2005).

42. Dronamraju, R. & Mason, J. M. Recognition of double strand breaks by a mutator protein (MU2) in Drosophila melanogaster. PLoS Genet. 5, e1000473 (2009).

43. Shao, Z. et al. Stabilization of chromatin structure by PRC1, a Polycomb complex. Cell 98, 37–46 (1999).

44. Francis, N. J., Saurin, A. J., Shao, Z. & Kingston, R. E. Reconstitution of a functional core polycomb repressive complex. Mol. Cell 8, 545–556 (2001).

45. Smith, E. R. et al. Drosophila UTX is a histone H3 Lys27 demethylase that colocalizes with the elongating form of RNA polymerase II. Mol. Cell. Biol. 28, 1041–1046 (2008).

46. Zou, L. & Elledge, S. J. Sensing DNA damage through ATRIP recognition of RPA-ssDNA complexes. Science 300, 1542–1548 (2003).

47. Zielke, N. et al. Fly-FUCCI: A versatile tool for studying cell proliferation in complex tissues. Cell Rep. 7, 588–598 (2014).

48. Brown, J. B. et al. Diversity and dynamics of the Drosophila transcriptome. Nature 512, 393–399 (2014).

49. LaRocque, J. R., Jaklevic, B., Su, T. T. & Sekelsky, J. Drosophila ATR in double-strand break repair. Genetics 175, 1023–1033 (2007).

50. Zhang, C. et al. Drosophila UTX coordinates with p53 to regulate ku80 expression in response to DNA damage. PLoS ONE 8, e78652 (2013).

51. Bordelet, H. & Dubrana, K. Keep moving and stay in a good shape to find your homologous recombination partner. Curr. Genet. 65, 29–39 (2019).

52. Saredi, G. et al. H4K20me0 marks post-replicative chromatin and recruits the TONSL– MMS22L DNA repair complex. Nature 534, 714–718 (2016).

53. Kallimasioti-Pazi, E. M. et al. Heterochromatin delays CRISPR-Cas9 mutagenesis but does not influence the outcome of mutagenic DNA repair. PLoS Biol. 16, e2005595 (2018).

54. Schep, R. et al. Impact of chromatin context on Cas9-induced DNA double-strand break repair pathway balance. Mol. Cell 81, 2216–2230.e10 (2021).

55. Jäger, N. et al. Hypermutation of the inactive X chromosome is a frequent event in cancer. Cell 155, 567–581 (2013).

56. Schulz, W. A., Lang, A., Koch, J. & Greife, A. The histone demethylase UTX/KDM6A in cancer: Progress and puzzles. Int. J. Cancer 145, 614–620 (2019).

57. Loubière, V. et al. Coordinate redeployment of PRC1 proteins suppresses tumor formation during Drosophila development. Nat. Genet. 48, 1436–1442 (2016).

58. O’Geen, H., Echipare, L. & Farnham, P. J. Using ChIP-seq technology to generate highresolution profiles of histone modifications. Methods Mol. Biol. 791, 265–286 (2011).

59. Dernburg, A. F. Formaldehyde fixation of Drosophila tissues onto slides for whole-mount FISH. Cold Spring Harb. Protoc. 2012, (2012).

60. Lerit, D. A., Plevock, K. M. & Rusan, N. M. Live imaging of Drosophila larval neuroblasts. J. Vis. Exp. (2014) doi:10.3791/51756.

