## Supplemental Figures for "Double-strand breaks in facultative heterochromatin require specific movements and chromatin changes for efficient repair"

### SUPPLEMENTAL FIGURE 1

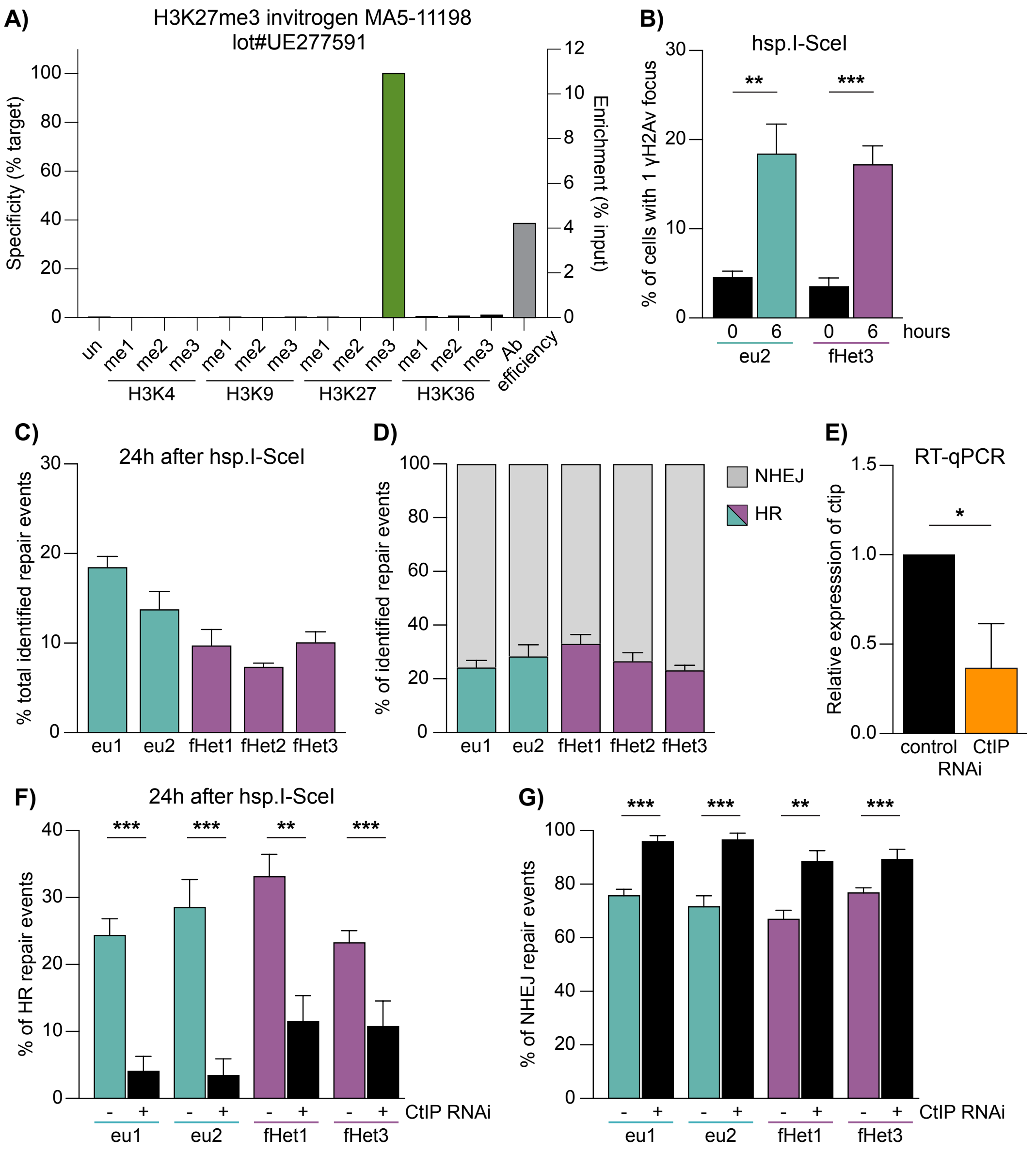


**Supplemental Figure S1 (extension Fig.1). DR-*white* system to induce single DSBs in euchromatin and facultative heterochromatin.**

**A**) ChIP specificity and efficiency of H3K27me3 antibody determined using EpiCypher K-MetStat panel. Bars indicate the specificity of the antibody for the expected target modification (green) and off-target modifications (black). Grey bars (right y-axis) indicate the percentage of target modification pulldown relative to input (measure of antibody efficiency). **B**) Wing discs of DR-*white*/hsp.I-SceI larvae were heat-shocked at 37°C for one hour to activate hsp.I-SceI. Number of DSBs per cell was quantified before heat-shock (0 hours) or after heat-shock (6 hours) using immunofluorescence staining for γH2Av (to visualize DSBs) and DAPI (to visualize nuclei). Bars indicate averages +SD of 3 wing discs per DR-*white* site (n≥160 cells per wing disc). **C, D**) DR-*white*/hsp.I-SceI larvae were heat-shocked for one hour to induce I-SceI. Repair products were subjected to PCR and Sanger sequencing 24 hours after I-SceI induction and analyzed using the TIDE algorithm. Graphs show quantification of total identified repair events (**C**) and the percentage of HR and NHEJ repair products (**D**). Bars indicate averages +SEM of ≥8 independent experiments (larvae) per condition. **E**) Relative CtIP expression level in 3rd instar larvae normalized to an internal control gene (vermillion), determined using Reverse Transcription followed by quantitative PCR (RT-qPCR). Bars indicate averages +SD of 4 single larvae per condition (luciferase control RNAi or CtIP RNAi). **F, G**) DR-*white*/hsp.I-SceI without (-) or with (+) CtIP RNAi were heat-shocked for one hour to induce I-SceI. Repair products were sequenced 24 hours after I-SceI induction and analyzed using the TIDE algorithm. Graphs show quantification of the percentage HR (**F**) and NHEJ repair products (**G**). Bars indicate averages +SEM of ≥5 independent experiments (larvae) per condition. (*) p-value≤0.05, (**) p-value≤0.01, (***) p-value≤0.001, unpaired t-test (**B, F, G**) and paired t-test (**E**).

### SUPPLEMENTAL FIGURE 2

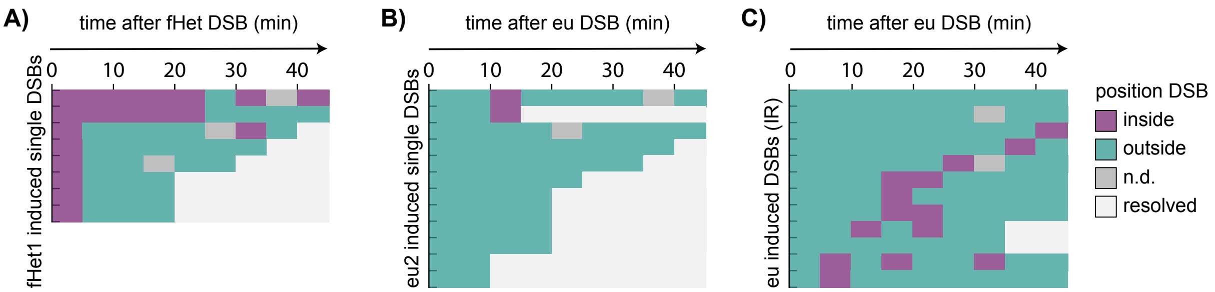


**Supplemental Figure S2 (extension Fig.2). Facultative heterochromatic DSBs move outside polycomb bodies.**

**A-C**) Quantification of live imaging of Mu2-eYFP (DSB) dynamics and kinetics relative to ph-p-mCherry domains. Rows indicate single Mu2-eYFP foci from the timepoint they appear (0 min) within (*fHet1* + ecDHFR-I-SceI (**A**)) or outside (*eu2* (**B**), 5Gy gamma-radiation (IR) (**C**)) polycomb bodies until the timepoint they resolve. Mu2-eYFP foci were followed up to 45 minutes after their appearance. Colors indicate localization of the Mu2-eYFP focus with respect to ph-p domains (inside polycomb body [purple], outside [green], not detectable [grey], or resolved [white]).

**SUPPLEMENTAL FIGURE 3**

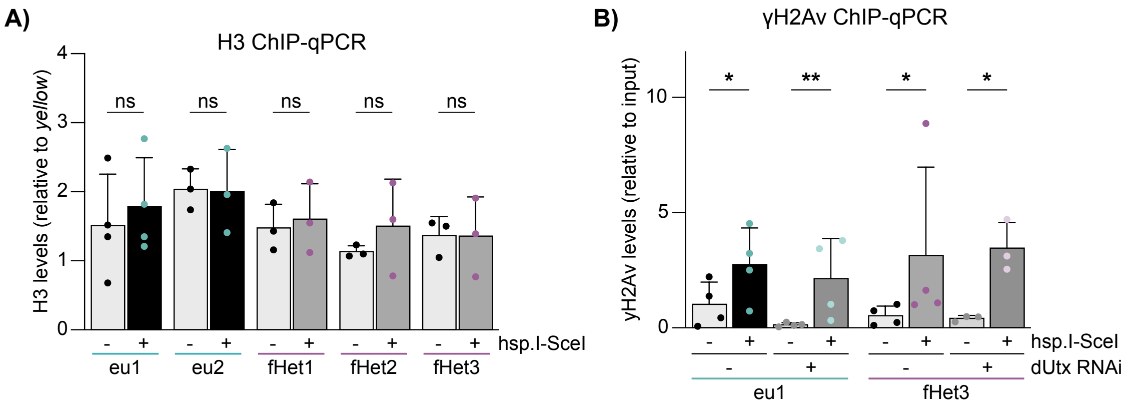


**Supplemental Figure S3 (extension Fig.3). DSBs in facultative heterochromatin do not locally change H3 or H3K27me1 levels.**

**A-B**) ChIP-qPCR analysis for H3 (**A**) and γH2Av (**B**) in the absence or presence of a DSB (-/+ hsp.I-SceI) in indicated conditions (as in Fig 1F). H3 levels were normalized using a *yellow* qPCR primer set as internal control. Averages are shown for ≥3 for independent experiments +SD. (ns) not significant, (*) p-value≤0.05, (**) p-value ≤0.01, paired t-test (**A**), ratio paired t-test (**B**).

### SUPPLEMENTAL FIGURE 4

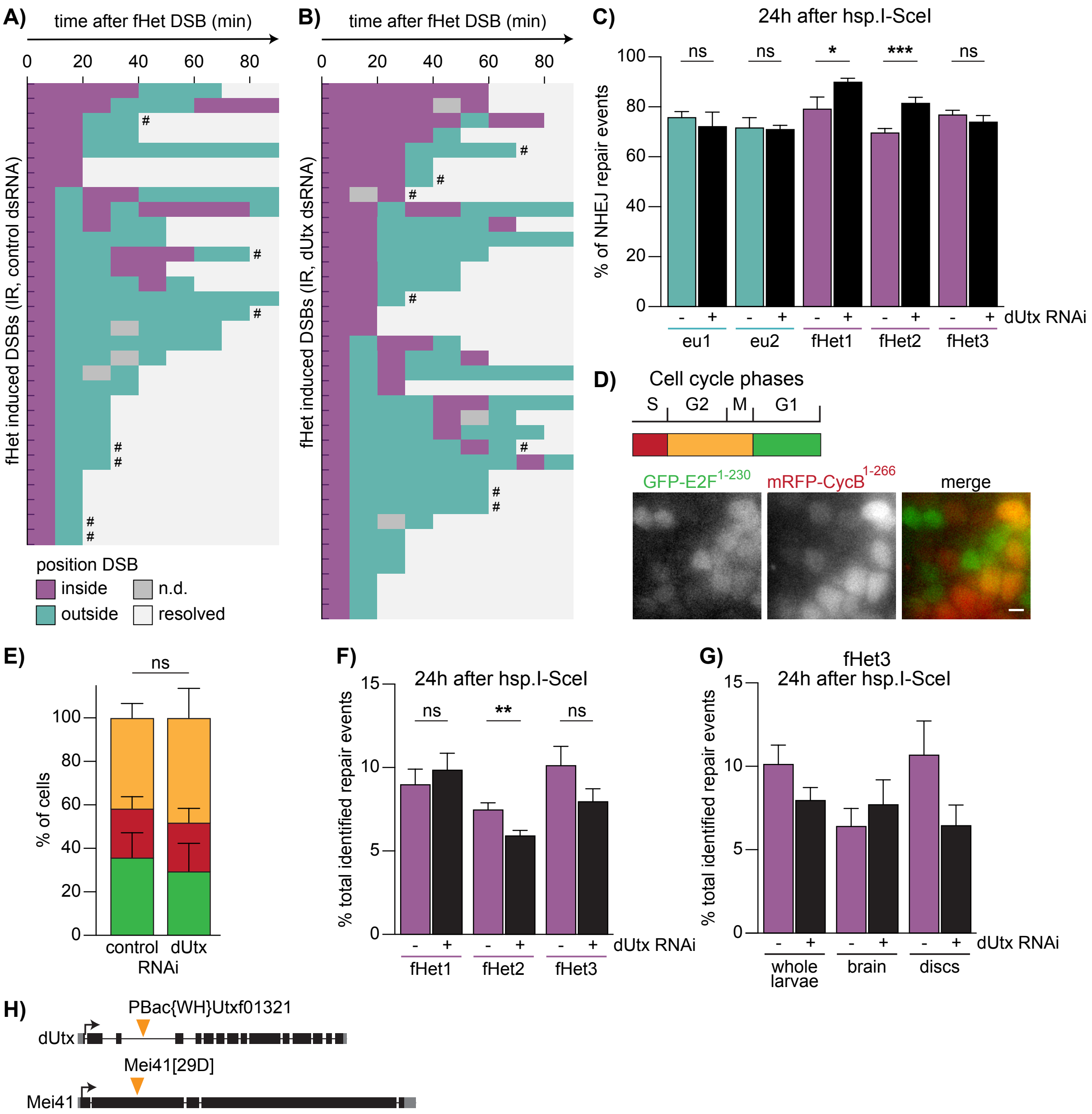


| Genotype | Viability | N |
|---|---|---|
| Mei41+; dUtx[f01321]/CyO | 0.6162 | 1534 |
| Mei41[29D]; dUtx[f01321]/CyO | 0.5163 | 2082 |
| Relative viability of double mutant: 84%<br>p-value: 0.0001 | | |

**Supplemental Figure S4 (extension Fig.4).**

**A, B**) Quantification of live imaging of Mu2-eYFP (DSB) dynamics and kinetics relative to the ph-p-mCherry domains upon 5Gy gamma-radiation (IR) to induce DSBs in control (**A**) or dUtx RNAi depleted (**B**) Kc cells. Each row indicates one single Mu2-eYFP focus that appeared within a polycomb body. Mu2-eYFP foci were followed up to 90 minutes after their appearance. Colors indicate localization of the Mu2-eYFP focus with respect to ph-p domains (inside polycomb body [purple], outside [green], not detectable [grey], or resolved [white]). # indicates DSBs that did not resolve during the course of our imaging experiment. **C**) DR-*white*/hsp.I-SceI with (+) or without (-) dUtx RNAi larvae were heat-shocked for one hour to induce I-SceI. Repair products were sequenced 24 hours after I-SceI induction and analyzed using the TIDE algorithm. Graph shows the percentage of identified NHEJ repair products. Bars indicate averages +SEM of ≥5 independent experiments (larvae). **D)** Top: Schematic of distinct colors indicating each cell cycle phase. Bottom: Representative images of the Fly-FUCCI system in wing discs. Error bar = 2µm. **E)** Cell cycle analysis in wing discs of control (luciferase RNAi) and dUtx depleted (dUtx RNAi) larvae using the Fly-FUCCI system as illustrated in D. Graph shows quantification of four discs per condition +SD (n≥400 cells per wing disc). (ns) not significant, unpaired t-test for the percentage of G1 cells. **F, G)** DR-*white*/hsp.I-SceI with (+) or without (-) dUtx RNAi larvae were heat-shocked for one hour to induce I-SceI. Repair products of whole larvae (F) or indicated tissues (G) were sequenced 24 hours after I-SceI induction and analyzed using the TIDE algorithm. Graph shows the percentage of total identified repair events per condition. Bars indicate averages +SEM of ≥5 independent experiments (larvae) (F) and averages +SD of ≥3 independent experiments (G)**. H**) Synthetic lethality assay of dUtx heterozygous mutant with ATR (mei41) mutant. N=1534 flies for the control line (dUtx mutant, wild-type ATR allele (mei41+; dUtx [f01321]/CyO) and n=2082 flies for the cross between dUtx- and ATR- mutant flies (mei41[29D]; dUtx[f01321]/CyO). (ns) not significant, (*) p-value≤0.05, (**) p-value≤0.01, (***) p-value≤0.001, unpaired t-test (**C, F, G**).
