## Supplemental Table for "Double-strand breaks in facultative heterochromatin require specific movements and chromatin changes for efficient repair"

| **Fly lines** | |
|---|---|
| *DR-white (eu 2)* | Janssen laboratory (Janssen et al., 2016) |
| *DR-white (others)* | |
| *ecDHFR-I-Sce* | Janssen laboratory (Janssen et al., 2016) |
| *hsp70.HA.I-Sce (hsp.I-Sce)* | Jan LaRocque (Do et al., 2014) |
| *mu2* -eYFP | Karpen laboratory (Janssen et al., 2016) |
| *php-mCh* | |
| *FlyFUCCI* | Bloomington #55123 |
| *UAS:dUtx RNAi* | VDRC 105986 |
| *UAS:ctip RNAi* | VDRC 100035 |
| *UAS:luciferase RNAi* | Bloomington #31603 |
| *Act5C-Gal4* | Bloomington #4414 and #3954 |

| **DR-white fly lines** | **Integration site** | **MiMIC line** | **BDSC #** |
|---|---|---|---|
| *DR-white eu2* | described in Janssen et al., 2016 | | |
| *DR-white eu3* | 3R (87A1) - 11876313 | MI00256 | 33599 |
| *DR-white fHet1* | 2R (60E3) - 24843209 | MI02312 | 58609 |
| *DR-white fHet2* | 3R (84A6) - 6866410 | MI06379 | 60795 |
| *DR-white fHet3* | 3R (90B5) -17590058 | MI02651 | 36022 |

| **qPCR primers** | **Goal** | **Fw** | **Rv** |
|---|---|---|---|
| *3xp3* | 1.4 kb away from I-SceI recognition site | CTCGCCCGGGGATCTAATTC | GCGACGTGTTCACTTTGCTT |
| *yellow* | internal control H3, yH2Av | ACGGTCCACAGAAGAGGATT | GCACTTAGCTCTAAGCTGACA |
| *ubx* | internal control H3K27me3 | ACAGAGGATTCCCTCTCTCG | CAGGCAGTCCTGTTTGTAGG |
| *fign1* | internal control H3K27me1 | CTAAAGGACTCACATGTCTC | GTAATCCCTCGTTTAGTCCT |
| *trx2* | internal control H3K27ac | AAAATTTAGCTGTGCTGCAA | CGAGTTACCCAATAGATGAC |
| *CtIP* | examine CtIP KD efficiency (larvae) | GAACTTGAAGCAAACTCGCC | CTTGACTGTGCTATTGCTG |
| *vermilion* | control for CtIP KD efficiency (larvae) | CTGCTCATGGACATCGACTC | GTGGACAGATTGAACAGATC |
| *dUtx* | examine dUtx KD efficiency (larvae + Kc cells) | CGGGGAGTTCCATATTGCCC | GGTACATCCATCCTAATTGTCGA |
| *tubulin* | control for CtIP KD efficiency (larvae) | GTGAAACACTTCCAATAAAAACTCAATATG | GCTCCAGTCTCGCTGAAGAA |

| **Primers** | **Goal** | **Fw** | **Rv** |
|---|---|---|---|
| *DR-white* | PCR primers repair product analysis | GGCCAGGGAACACCTGATTT | CGCGAATTCGTCGACATAAC |
| *DR-white sequencing* | sequencing primer repair product analysis | GAGCCCACCTCCGGACTGGAC | - |
| *dsRNA dUtx* | induce dUtx KD in Kc cells | TAATACGACTCACTATAGgcaacttcaacgccatgcc | TAATACGACTCACTATAGGgcgcattcacttgcaaaatgg |
| *dsRNA yellow* | induce control KD in Kc cells | TAATACGACTCACTATAGggaaaaactaagccaacgtcatc | TAATACGACTCACTATAGGgccgtggatataggcaaaaa |
